# Baculovirus *bv/odv-e26* is required for host behavioral manipulation by optimizing viral virulence to lepidopteran hosts

**DOI:** 10.1101/2022.05.26.493567

**Authors:** Hiroyuki Hikida, Susumu Katsuma

## Abstract

Host behavioral manipulation is a widely used strategy for parasites to enhance their transmission. Lepidopteran nucleopolyhedroviruses (NPVs) induce hyperactivity and alteration of behavioral patterns in host insect larvae. Previous studies revealed that *Bombyx mori nucleopolyhedrovirus* (BmNPV) does not induce hyperactivity in larval *Bombyx mori*, when *Bm8*, a homologue of the baculovirus *bv/odv-e26* gene, is knocked out. *Bm8* knockout also enhances pathogenicity, but the relationship between *Bm8* functions and host behavioral manipulation is still unknown. This study revealed that *Bm8* is dispensable for altering behavioral patterns but crucial for inducing hyperactivity. *Bm8* overexpression decreased pathogenicity in *B. mori* larvae and delayed viral infection in cultured cells. These results suggest that *Bm8* maintains larval locomotory activity and synchronizes hyperactivity induction with behavioral pattern alteration by suppressing viral virulence. Conserved amino acid residues were also identified in the coiled-coil domain of the Bm8 protein, which is required for suppressing viral virulence in cultured cells and larval hosts. Although *bv/odv-e26* is previously thought to be conserved in a limited group of lepidopteran NPVs, our synteny-based bioinformatics approach discovered putative homologues in the genomes of most lepidopteran NPVs. Functional analyses revealed that phylogenetically close and distant *bv/odv-e26* homologues possess suppressive activity with functionally essential common residues in their coiled-coil domains. Collectively, these findings indicate that baculovirus *bv/odv-e26* is a conserved gene optimizing viral virulence for establishing behavioral manipulation in NPV-infected larval hosts.

**Author Summary:** Baculovirus-induced host abnormal behavior has been described in various lepidopteran insects for a long time, although it is still largely unknown how the virus establishes this complex trait. We showed that baculovirus *bv/odv-e26* is a factor that suppresses viral virulence and synchronizes hyperactivity and behavioral pattern alteration. We also revealed that putative *bv/odv-e26* homologues are encoded by a wider range of baculoviruses than previously thought, with functionally conserved residues. We confirmed the suppressive function in both phylogenetically close and distant *bv/odv-e26* homologues. These results indicate that acquiring a suppressive factor for viral virulence is a crucial event for the evolution of a host manipulation strategy by baculoviruses.

## Introduction

Alteration of host behavior by parasitic infections is widely observed in the environment. Some parasites manipulate their host behavior to efficiently transmit their infection [1]. Baculovirus is a pathogen manipulating host caterpillars’ behavior to spread its infection. Baculovirus-infected caterpillars exhibit vigorous movement at a late stage of infection and are finally led to the elevated position, where they die [2, 3]. Host death at the elevated position is thought to facilitate viral spreading by rainfall or avian feeding [4, 5]. Some baculoviruses also encode cysteine protease and chitinase that efficiently degrade the host insect’s body and enhance progeny virus release into the environment [6, 7]. Moreover, baculoviruses produce protein crystals, called occlusion bodies (OBs), which protect virions from environmental stresses and maintain their infectivity for an extended period [8].

The family *Baculoviridae* is a group of insect-specific large double-stranded DNA viruses of 100 to 180 kilobase pair (kbp) genome with rod-shaped, enveloped virions. The family is phylogenetically divided into four genera: *Alphabaculovirus, Betabaculovirus, Gammabaculovirus*, and *Deltabaculovirus* [9–11]. *Alphabaculovirus*, also known as lepidopteran nucleopolyhedrovirus (NPVs), is further divided into groups I and II [12]. Baculoviruses encode 80 to 180 genes that are coordinately expressed to optimize their infection cycle [13]. They are clustered into four types based on their expression timing. After viral entry, immediate-early genes are expressed and initiate virus infection. Delayed-early genes are expressed with the support of some immediate-early genes and facilitate viral genome replication and expression of late genes [14]. Late genes are transcribed by viral RNA polymerase and encode proteins that composes nucleocapsids [15–17]. A part of preceding genes supports the expression of very late genes, *polh* and *p10* [18]. These two genes encode extraordinarily highly expressed proteins that composes OBs and facilitates their release into the extracellular environment, respectively [8, 19].

Host behavioral manipulation has been observed in alphabaculoviruses and comprises a combination of hyperactivity and vertical movement [2, 3, 20–22]. Previous studies identified virus-encoded *protein tyrosine phosphatase (ptp)* and *ecdysteroid UDP-glucosyltransferase (egt)*, which are required for the sufficient upregulation of horizontal and vertical activities, respectively, in several host–virus combinations [20–23]. Succeeding studies suggested that, without these genes, viruses fail to establish their infection in crucial tissues for host behavioral control at the appropriate timing, disrupting the induction timing of host behavioral manipulation [23, 24]. A similar mechanism was suggested in the case of deletion of Bombyx mori nucleopolyhedrovirus (BmNPV) *actin rearrangement factor 1 (arif-1)* and *Bm96*; both alter larval survival time and mitigate larval locomotory activity at a late stage of infection [25, 26].

The baculovirus *bv/odv-e26* gene was originally found in a genomic library of Autographa californica multiple nucleopolyhedrovirus (AcMNPV), a prototype of alphabaculoviruses, and suggested to be implicated in late gene expression [17]. The encoded protein harbors a coiled-coil domain important for protein interaction and interacts with viral *trans*-activator IE1 [27]. Phylogenetic analysis indicated that this gene is found only in the genomes of group I alphabaculoviruses [12, 28]. BmNPV, a close relative of AcMNPV, encodes a homologous protein called Bm8, which colocalizes with IE1 and genomic enhancer region *homologous regions (hrs)* [29, 30]. This complex exhibits punctate localization in the cell nucleus, suggesting that Bm8 is also involved in the regulation of gene expression. In infected *B. mori* larvae, *Bm8* deletion from the BmNPV genome results in a fast-killing phenotype and lowers host locomotory activity at a late stage of infection [31, 32]. *Bm8*-deleted BmNPVs also exhibit abnormal tissue tropism in multiple tissues, particularly remarkable in larval middle silk glands (MSGs) [31, 33]. As *Bm8* deletion results in a fast-killing phenotype and enhances virulence, it was suggested that its deletion disrupts the induction timing of host behavioral manipulation [31].

A new recording system that enables the continuous observation of the infected larvae was recently developed. The method provides high-resolution data for baculovirus-induced host behavioral manipulation, revealing that the abnormal behavior of BmNPV-infected *B. mori* larvae comprises hyperactivity and alteration of behavioral patterns [34]. In this study, locomotory phenotypes of *B. mori* larvae infected with *Bm8*-deleted BmNPV were thoroughly investigated with this system. Combined with the pathogenic properties of *Bm8* mutants in *B. mori* larvae and cultured cells, it was concluded that *Bm8* optimizes viral virulence to larval hosts for synchronizing hyperactivity and behavioral pattern alteration. This study also identified orthologous genes of *bv/odv-e26* from group II alphabaculoviruses. Mutagenesis experiments revealed the functional importance of two conserved residues in the coiled-coil domain of the Bm8 protein, which was also observed in an orthologous protein from phylogenetically distant alphabaculoviruses. Taken together, these findings indicate that *bv/odv-e26* orthologues are commonly utilized to optimize viral virulence for establishing behavioral manipulation in alphabaculovirus-infected larvae.

## Results

### *Bm8* is dispensable for locomotory pattern alteration but required for hyperactivity in BmNPV-infected *B. mori* larvae

Locomotory patterns of *B. mori* larvae infected with wild-type (T3) and *Bm8*-deleted (Bm8D) BmNPVs were compared to those of mock-infected larvae. As described previously [34, 35], mock-infected larvae periodically repeated stational and active behaviors, whereas T3-infected larvae showed continuous locomotion at a late stage of infection (Fig. 1A; Fig. S1). This periodical pattern in mock-infected larvae was lost in those infected with Bm8D, and locomotion of Bm8D-infected larvae was continuously induced at ~80 to 90 hours post infection (hpi) (Fig. 1A; Fig. S1). These results indicated that the locomotory pattern is also disrupted in Bm8D-infected larvae. In contrast, the locomotion of Bm8D-infected larvae was frequently intermitted, and their locomotory activity was less enhanced, compared to those of T3-infected larvae. Consistent with these results, Bm8D-infected larvae showed a lower travel distance than T3-infected larvae, which is almost the same level as that of mock-infected larvae (Fig. 1B). The median locomotory speed and duration of Bm8D-infected larvae were also comparable to those of mock-infected larvae and significantly lower than those of T3-infected larvae (Fig. 1C and D). These results demonstrated that Bm8D fails to induce hyperactivity at a late stage of infection.

**Fig 1.**
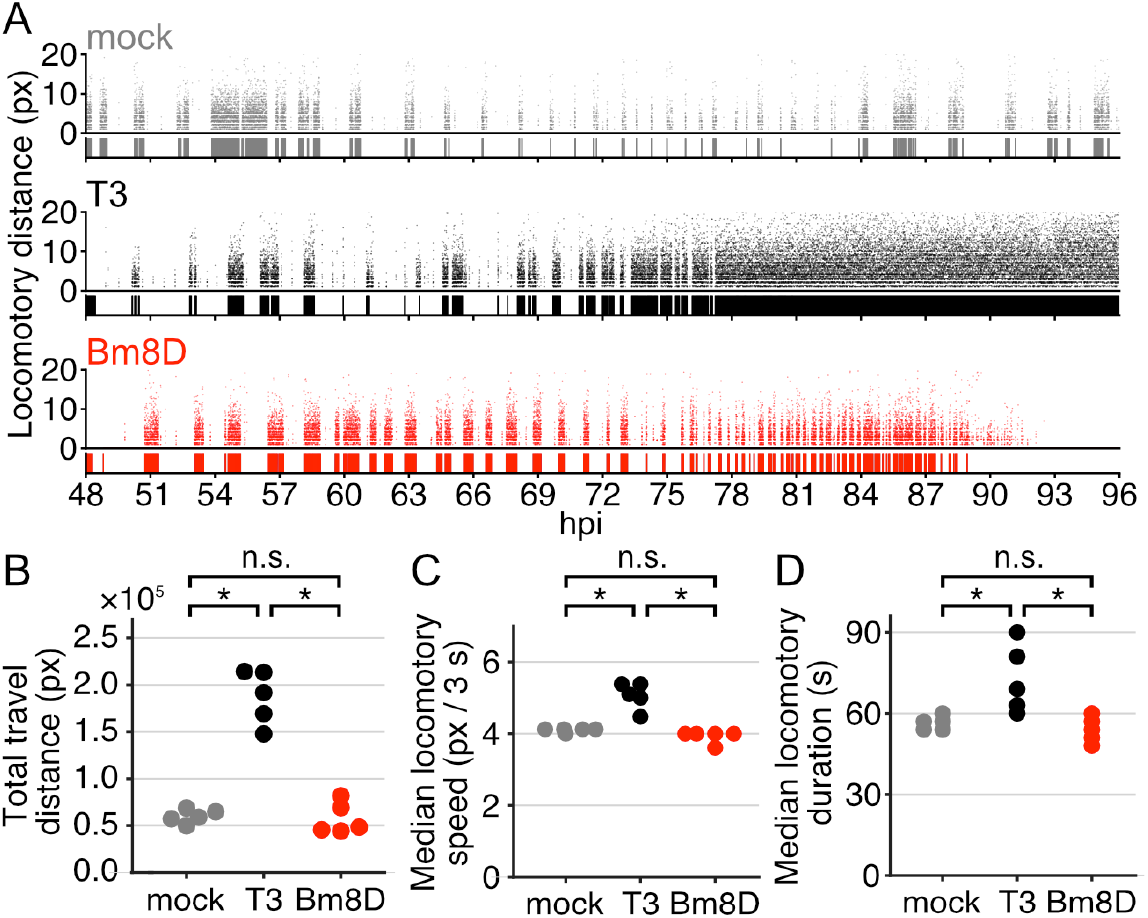
Locomotory activity of mock-infected (gray), T3-infected (black), and Bm8D-infected (red) larvae. (A) Temporal pattern of locomotory activity of infected larvae. The X-axis shows hours post infection (hpi), and the Y-axis shows locomotory distance at 3 s intervals. The diagram below each scatter plot indicates the duration of larval locomotion. Results of other individuals are shown in Fig. S1. (B) Total travel distance of infected larvae. (C) Median locomotory speed of infected larvae. (D) Median locomotory duration of infected larvae. (B–D) Each point indicates an individual larva (n = 5). (\**p*< 0.05, log-rank test followed by Bonferroni correction. n.s., no significant difference).

### *Bm8* delays viral infection in both *B. mori* larvae and cultured cells

Previous studies revealed that *Bm8* deletion enhances viral virulence [31, 33]. This study showed that *Bm8* is required for hyperactivity but dispensable for behavioral pattern alteration (Fig. 1). These results provided a hypothesis that *Bm8* inhibits the progression of viral infection and maintains larval locomotory activity until the behavioral pattern is altered. To examine this hypothesis, recombinant BmNPV that overexpresses *Bm8* (Bm8OE; Fig. 2A) was generated, and its phenotypes were investigated in *B. mori* larvae. Bm8OE significantly extended host survival time, whereas Bm8D showed a significant fast-killing phenotype compared to T3 (Fig. 2B). All larvae infected with Bm8OE died, indicating that *Bm8* delays the progression of viral infection but does not completely inhibit it. In some alphabaculoviruses, including BmNPV, viral cathepsin (V-CATH) is crucial for viral virulence and host survival time [36]. Therefore, V-CATH accumulation was examined in hemolymph and fat bodies of infected larvae. V-CATH expression and accumulation of Bm8OE-infected larvae were lower in fat bodies and hemolymph than those infected with T3 and Bm8D, and mature V-CATH was not detected in Bm8OE-infected larvae (Fig. 2C). V-CATH activity was not detected in the hemolymph of Bm8OE-infected larvae, whereas the hemolymph of T3- and Bm8D-infected larvae showed slight and high V-CATH activity, respectively (Fig. 2D). These results demonstrated that *Bm8* suppresses viral virulence and delays the progression of viral infection in *B. mori* larvae.

**Fig 2.**
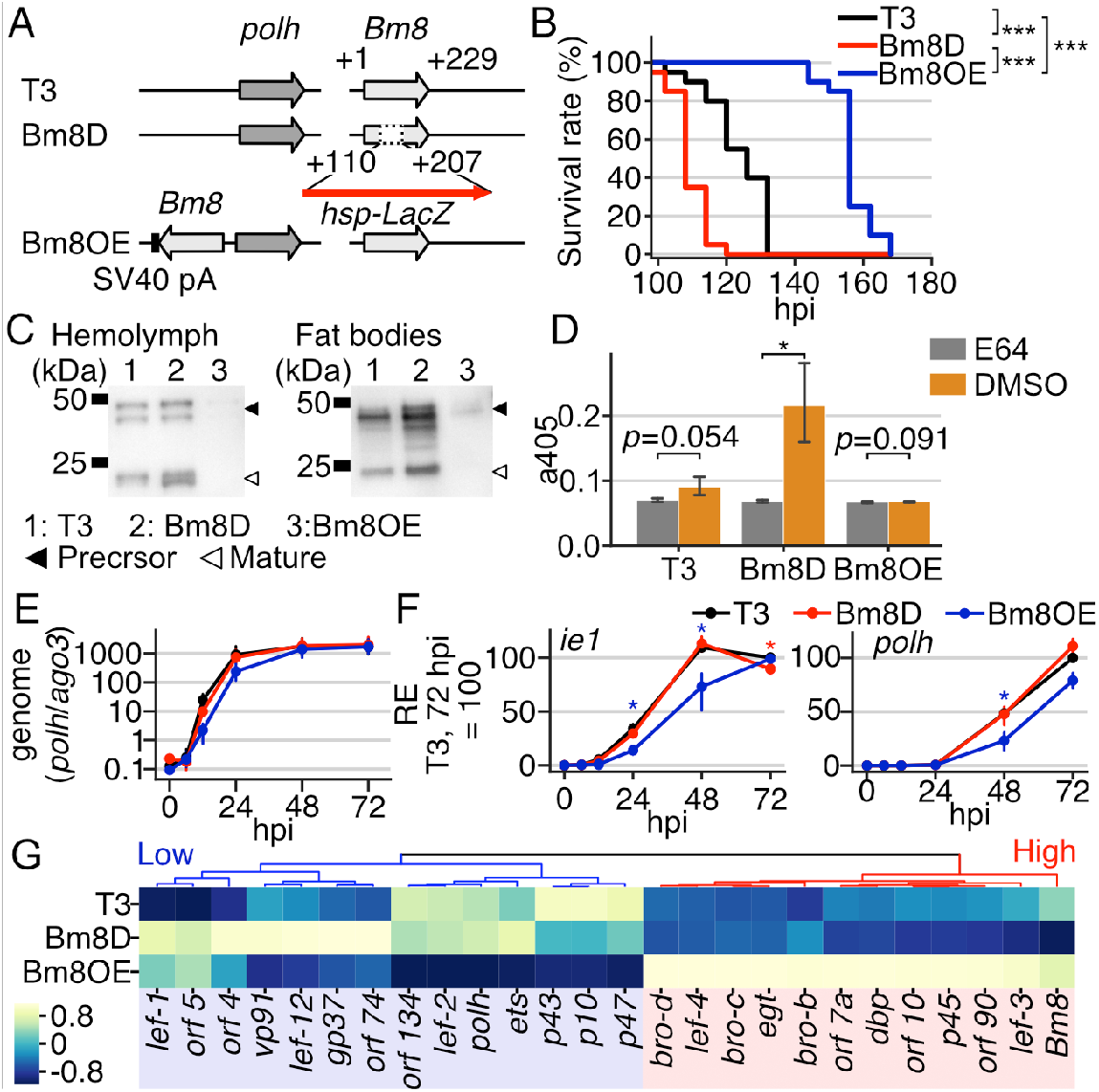
Generation of a Bm8-overexpressing BmNPV and its phenotypes in infected *B. mori* larvae and cells. (A) Schematic image of recombinant viruses. (B) Survival curves of infected larvae (n = 20, \*\*\**p*< 0.005, pairwise log-rank test). LT_50_ was 126, 108, and 156 h for T3-, Bm8D-, and Bm8OE-infected larvae, respectively. (C) V-CATH accumulation and expression in larval hemolymph and fat bodies at 90 hpi. (D) V-CATH activity in larval hemolymph (n = 3). E64 was used as a cysteine protease inhibitor. (\**p*< 0.05, one-side paired-samples *t*-test). (E) Viral genome replication in infected cells (n = 3). (F) Temporal expression of *ie1* and *polh.* Blue and red asterisks indicate that Bm8OE and Bm8D show significant differences from the other two viruses, respectively. *p*< 0.05, Tukey’s HSD test. (D–F) Error bars indicate 95% confidence intervals. (G) Clustered heatmap of viral gene expression at 48 hpi. Genes showing a significantly different expression with more than twofold change are shown (n = 2). Blue and red indicate the clusters in which the genes show low and high expression in Bm8OE-infected cells, respectively. The results of the third replicate are shown in Fig. S3.

Bm8OE phenotypes were further investigated in *B. mori* cultured cells. The viral genome replication patterns were similar between Bm8D and T3 (Fig. 2E). In contrast, the viral replication level in Bm8OE-infected cells was slightly lower at an early stage of infection and became similar to other viruses at a late stage. Also, expression levels of *ie1* and *polh*, immediate-early and very late genes, respectively, were significantly lower in Bm8OE-infected cells than those of the other two viruses (Fig. 2F). Similar expression patterns were observed in *lef-2* and *vp39*, delayed-early and late genes, respectively (Fig. S2). The global expression pattern of viral genes was further investigated by RNA sequencing at 48 hpi (Fig. 2G; Fig. S3). Genes with differential expression patterns were largely classified into two groups by expression levels in Bm8OE-infected cells. Highly expressed genes in Bm8OE-infected cells included early genes (i.e., *bro* genes and *egt*), whereas those with low expression included very late genes, *polh* and *p10*. These results demonstrated that *Bm8* delays the progression of viral infection in host cells, leading to prolonged viral infection in host larvae.

### Conserved amino acid residues in the coiled-coil domain are required for the suppressive function of the Bm8 protein

The coiled-coil domain of the Bm8 protein is required for the interaction with IE1 [30], suggesting that the domain is important for *Bm8* to suppress viral infection. Three recombinant BmNPVs were generated, in which a point mutation was introduced at an amino acid residue conserved in all homologues of the Bm8 protein found in group I alphabaculoviruses (Fig. 3A and B; Fig. S4). These recombinant BmNPVs were inoculated to *B. mori* larvae, and OB formation in MSGs was investigated (Fig. 3C). As reported previously [31, 33], few OBs were observed in MSGs of T3-infected larvae, whereas Bm8D produced many OBs in MSGs. The introduction of an amino acid substitution at the 88^th^ leucine did not affect the OB production level in MSGs. In contrast, mutations in the 96^th^ isoleucine or 99^th^ leucine resulted in mass OB production, similar to the Bm8D phenotype. These results indicated that two residues, the 96^th^ isoleucine and 99^th^ leucine, in the coiled-coil domain are required for the suppressive function of the Bm8 protein in *B. mori* larvae.

**Fig 3.**
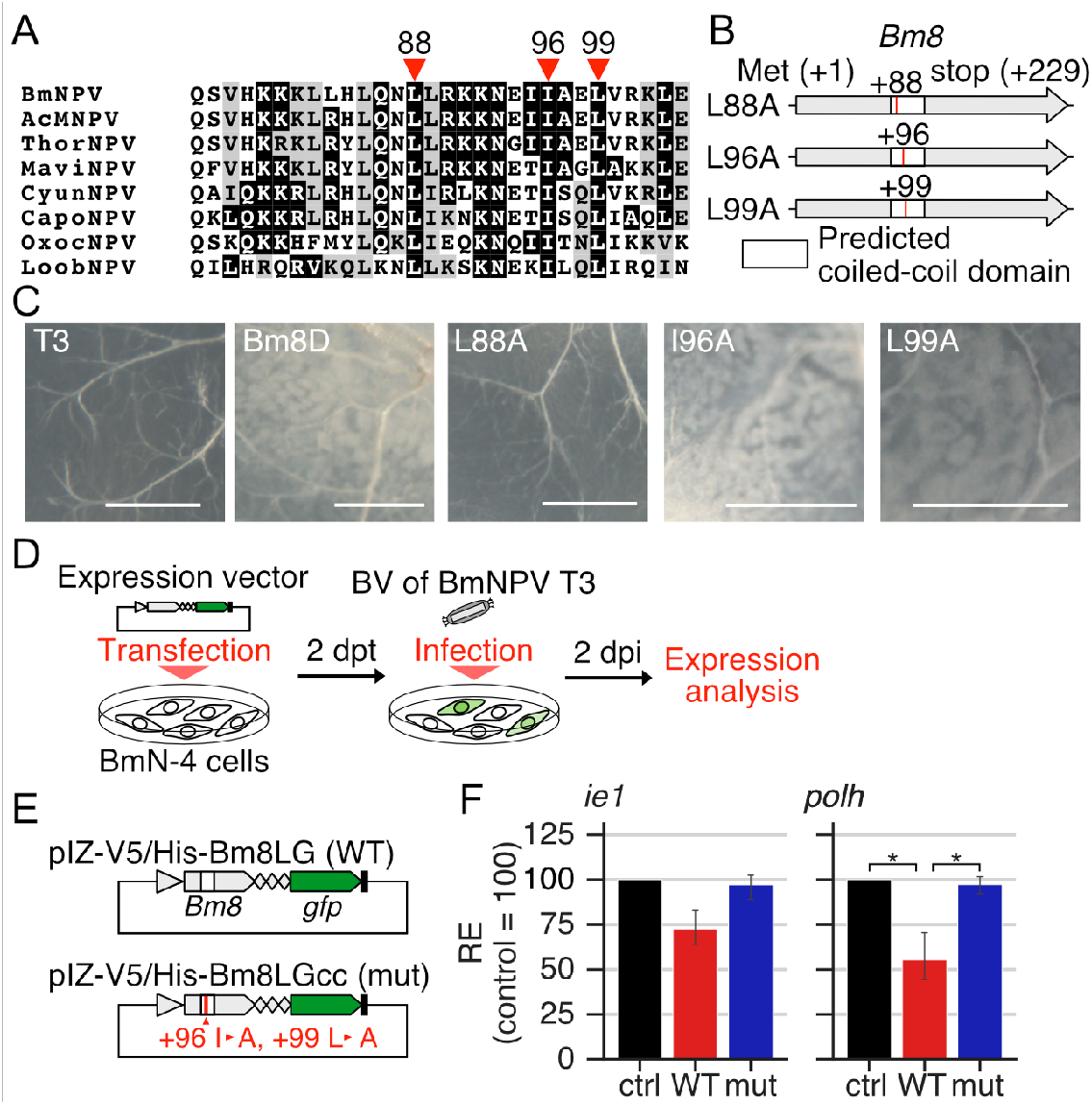
Identification of the conserved residues required for the Bm8 protein function. (A) Selected alignment of the coiled-coil domains of Bm8 homologues. Red arrowheads indicate residues conserved in all homologues. The numbers of the residues in the Bm8 protein are indicated above the arrowheads. The abbreviations for viruses are listed in Table S2. The alignment of all homologues is shown in Fig. S4. (B) Schematic image of point mutations introduced in recombinant viruses. Red lines indicate the position of point mutations in the coiled-coil domain. (C) Microscopic images of infected MSGs. White blobs are OB-containing cells. Bars, 200 μm. Schematic images of (D) experimental procedure and (E) plasmid vectors for (F). (E) The red line indicates point mutations in the coiled-coil domain. (F) Relative expression of *ie1* and *polh* in plasmid-transfected cells at 48 hpi. The expression level in cells transfected with the control vector (pIZ-V5/His-GFP) is set as 100. Error bars indicate 95% confidence intervals (n = 3). (\**q*< 0.05, paired-samples *t*-test followed by Benjamini–Hochberg correction.)

The importance of the 96^th^ isoleucine and 99^th^ leucine in the Bm8 protein function was further investigated using *B. mori* cultured cells. Plasmid vectors expressing green fluorescent protein (GFP)-fused wild-type Bm8 or GFP-fused Bm8 derivative with the 96^th^ alanine and 99^th^ alanine were constructed and transfected into BmN-4 cells, followed by T3 infection (Fig. 3D and E). GFP fluorescence in cells transfected with the wild-type vector was observed uniformly in the nucleus without virus infection and changed to punctate loci in the nucleus after infection (Fig. S5). In contrast, transfection of the mutant vector resulted in weak and almost no fluorescence before and after infection, respectively. *polh* expression in cells transfected with the wild-type vector was significantly lower than in control cells, and this decrease was not observed in mutant vector-transfected cells (Fig.3F). Similar trends were obtained in *ie1* expression, although the differences were not significant. These results showed that the 96^th^ isoleucine and 99^th^ leucine are required for the suppressive function of the Bm8 protein in *B. mori* cultured cells.

### *bv/odv-e26* and its suppressive function are widely conserved in alphabaculoviruses

Previous and present results strongly suggested that the inhibitory role of *Bm8* for viral infection is crucial for the appropriate induction of host abnormal behavior in BmNPV. In contrast, *bv/odv-e26* homologues are thought to be conserved only in group I alphabaculoviruses, although the host behavioral manipulation has been described in group II alphabaculoviruses [2, 3, 12, 20, 23, 28]. Some group II alphabaculoviruses have genes located between *egt* and *Bm9* homologues, whose product sizes are similar to that of *bv/odv-e26* [37]. As all sequenced alphabaculoviruses harbor *egt* and a large number of them encode *Bm9* homologues, it was speculated that phylogenetically distant *bv/odv-e26* homologues are located between the two genes in some genomes of group II alphabaculoviruses. To examine the hypothesis, all open reading frames (ORFs) located between *egt* and *Bm9* homologues were selected, and their homologous proteins were searched for alphabaculovirus genomes registered as a reference sequence in the National Center for Biotechnology Information (NCBI) GenBank database (Fig. S6). The results revealed that most alphabaculoviruses harbor a homologue of *bv/odv-e26* between *egt* and *Bm9* (Fig. 4A). Some alphabaculoviruses did not possess *Bm9* homologues but harbored *bv/odv-e26* homologues downstream of *egt*. Among 54 alphabaculovirus genomes obtained from the NCBI RefSeq database, *bv/odv-e26* homologue was not detected only in the genome of Operophtera brumata nucleopolyhedrovirus (OpbuNPV), which is the most ancient type of alphabaculovirus [38].

**Fig 4.**
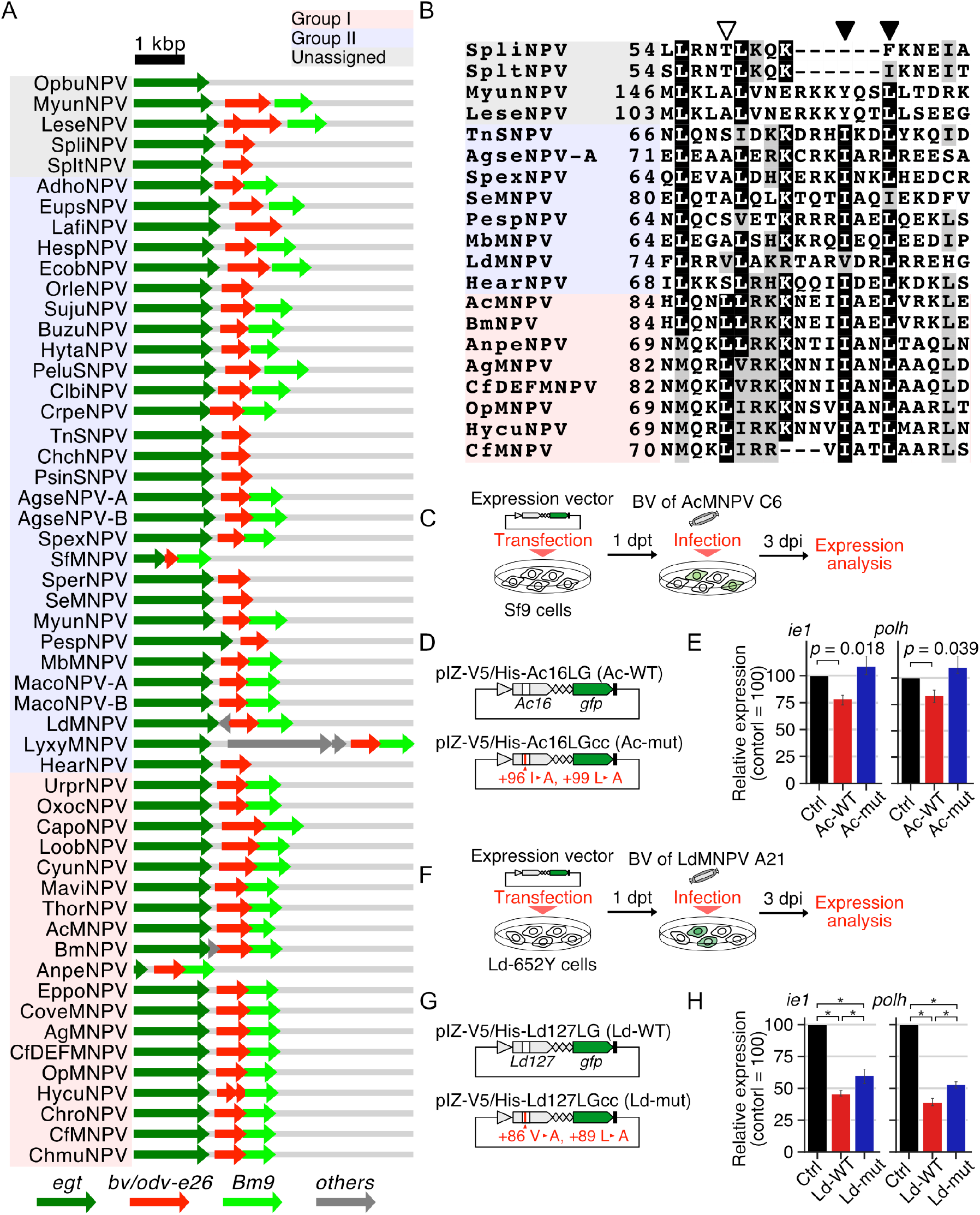
Identification and characterization of *bv/odv-e26* homologues. (A) Distribution of *bv/odv-e26* homologues in alphabaculovirus genomes. (B) Alignment of coiled-coil domains. The white arrowhead indicates Bm8 88^th^ residue. Black arrowheads indicate 96^th^ and 99^th^ residues. (A and B) The red, blue, and gray background colors indicate groups I and II and unassigned alphabaculoviruses, respectively. Schematic images of (C) experimental procedure and (D) plasmid vectors for (E). (E) Relative expression of *ie1* and *polh* in plasmid-transfected Sf-9 cells after AcMNPV infection. *p*-values were calculated by paired-samples *t*-test and corrected by Benjamini-Hochberg correction but no significant difference was detected after the correction. Schematic images of (F) experimental procedure and (G) plasmid vectors for (H). (H) Relative expression of *ie1* and *polh* in plasmid-transfected LdMNPV-infected Ld652Y cells. (\**q* < 0.05, paired-samples *t*-test followed by Benjamini-Hochberg correction). (D and G) The red line indicates point mutations in the coiled-coil domain.

Deduced amino acid sequences of *bv/odv-e26* homologues were aligned, and their phylogenetic tree was constructed. The tree topology was largely consistent with that constructed by 38 baculovirus core genes, and homologues in group I and II alphabaculoviruses were phylogenetically separated (Fig. S7). The alignment of deduced amino acid sequences of these homologues showed the presence of highly conserved coiled-coil domains (Fig. 4B; Fig. S8). Particularly, the amino acid residues 96^th^ isoleucine and 99^th^ leucine, essential for the suppressive function of the Bm8 protein, were conserved in most groups I and II alphabaculoviruses. In some group II alphabaculoviruses, the corresponding residues were replaced with structurally similar amino acids (i.e., valine, leucine, or isoleucine). The residue 88^th^ leucine, dispensable for Bm8 protein function (Fig. 3B and C) was not conserved in group II alphabaculoviruses. These results suggested that *bv/odv-e26* homologues have similar suppressive functions.

The functions of *bv/odv-e26* homologues were investigated using AcMNPV *Ac16* and LdMNPV *Ld127*, which are close and distant relatives of *Bm8*, respectively. GFP-fused wild-type or mutant proteins were transiently expressed in susceptible cell lines for each virus (Fig. 4C and F). In mutant proteins, two amino acid residues corresponding to the 96^th^ isoleucine and 99^th^ leucine of the Bm8 protein were replaced with alanine (Fig. 4D and G). In AcMNPV, wild-type Ac16 suppressed *ie1* and *polh* expression compared to control (*p*= 0.018 and 0.039, respectively), whereas the mutant protein almost lost this suppressive activity (*p*= 0.23 and 0.19, respectively; Fig. 4E). LdMNPV Ld127 significantly suppressed *ie1* and *polh* expression; unlike group I homologues, mutant Ld127 still had a suppressive activity that was significantly weaker than that of wild-type (Fig. 4H). Wild-type Ac16 and Ld127 showed punctate localization in the host cell nucleus (Fig. S9). Like the Bm8 protein, the mutant Ac16 protein showed no fluorescence, but the mutant Ld127 protein maintained GFP fluorescence after virus infection. These results indicated that *bv/odv-e26* orthologues are conserved in most alphabaculoviruses with their suppressive activity, but the proteins encoded by groups I and II alphabaculoviruses possess different biochemical properties.

## Discussion

Baculovirus-induced host behavioral manipulation has been described for a long time, but its underlying mechanisms are largely unknown. A recent study showed that the abnormal behavior of larval *B. mori* is composed of two sequential events: enhancement of locomotory activity and drastic alteration of the locomotory pattern [34]. This study focused on a baculovirus gene *bv/odv-e26*, and its role in host behavioral manipulation. *Bm8*, a BmNPV homologue of *bv/odv-e26*, is dispensable for behavioral pattern alteration, but its deletion results in lower locomotory activity comparable to that of mock-infected larvae (Fig. 1; Fig. S1). Combined with the finding that *Bm8*-deleted BmNPVs showed a fast-killing phenotype [31, 32], it was suggested that *Bm8* inhibits viral infection, and its deletion impairs larval locomotory activity when the behavioral pattern was altered. Using a *Bm8*-overexpressing virus, we showed that *Bm8* suppresses and delays virus infection in both *B. mori* cultured cells and larvae (Fig. 2). Collectively, these results demonstrated that *Bm8* optimizes viral virulence and synchronizes hyperactivity induction with behavioral pattern alteration, essentially contributing to the successful manipulation of host behavior.

*bv/odv-e26* homologues are thought to be found only in group I alphabaculoviruses [12, 28]. However, group II alphabaculoviruses also induce abnormal behavior in host insects. In this study, we found that the homologous sequences of *bv/odv-e26* are widely distributed in alphabaculovirus genomes (Fig. 4A). The homologues were found downstream of the *egt* locus, and the encoded proteins harbor conserved coiled-coil domains (Fig. 4B). Particularly, essential residues for the suppressive function of the Bm8 protein were well conserved in group II alphabaculoviruses (Figs. 3 and 4B). In addition, AcMNPV Ac16 and LdMNPV Ld127, which are phylogenetically close and distant homologues of the Bm8 protein, respectively, suppress viral gene expression in their susceptible cell lines (Fig. 4C–H). These results indicated that *bv/odv-e26* homologues in a wide range of alphabaculoviruses are orthologues with conserved functions. Among all alphabaculovirus genomes registered in the NCBI RefSeq database, only OpbuNPV does not encode a homologue of *bv/odv-e26* (Fig. 4A). As OpbuNPV is thought to be the most ancestral type of alphabaculovirus [38], it is suggested that *bv/odv-e26* was acquired at an early time in the course of alphabaculovirus evolution. Because there is no literature to refer to concerning OpbuNPV-induced abnormal behavior in host insects, it cannot be concluded that *bv/odv-e26* is a mandatory factor for host behavioral manipulation by all alphabaculoviruses. Nevertheless, these results indicated that *bv/odv-e26*-mediated optimization of viral virulence is crucial for establishing host behavioral manipulation in alphabaculoviruses.

Transiently expressed GFP-fused Bm8 protein with point mutations in the coiled-coil domain showed a faint GFP fluorescence at 2 dpi, whereas GFP-fused wild-type Bm8 remained at a visible level at the same time point (Fig. S5). Previous study showed that the Bm8 protein level peaks at ~24 hpi and reduces rapidly at a late stage of infection [31]. These results suggested that Bm8 is actively degraded, and the coiled-coil domain mediates protein stability. This temporal change in Bm8 may contribute to the precise regulation of viral virulence through a stage-dependent suppressive activity. Similar dynamics were found in the AcMNPV homologue, the Ac16 protein, whereas the mutant Ld127 protein was stable and retained fluorescence in the nuclei as observed in wild-type Ld127 (Fig. S9). Consistent with this observation, the mutant Ld127 protein also had suppressive activity, whereas Bm8 and Ac16 mutants did not (Figs. 3F and 4E and H). These results suggested that BV/ODV-E26 exhibits different biochemical properties during infection between groups I and II viruses. One of the large differences between the groups is the membrane fusion protein for cell entry. Group I alphabaculoviruses utilize GP64, whereas group II viruses use another fusion protein called F, thought to be displaced by GP64 in group I viruses [8, 12, 39–41]. As this study indicated that BV/ODV-E26 aims to optimize viral virulence in infected larvae, viral spread pattern is crucial for determining its biochemical properties. Collectively, it can be speculated that the functional difference of the BV/ODV-E26 proteins between groups I and II resulted from the difference in the cell-entry manner of these groups. This difference may also be related to the divergence of protein sequences, which hampered the identification of its homologues in group II viruses.

This study showed that *bv/odv-e26*, a suppressive factor for viral virulence, is required for sufficient locomotory activity and conserved in a wide range of baculoviruses. Previously, *ptp* and *arif-1* were shown to enhance viral virulence and larval locomotory activity at a late stage of infection [24, 25]. Taken together, these results indicated that positive and negative regulation of viral virulence is crucial for successful host behavioral manipulation in baculoviruses. Currently, the infection dynamics of baculovirus in larval tissues are not fully understood. As *bv/odv-e26* is also involved in tissue tropism determination [31, 33], further studies of this gene might highlight the suppression of viral virulence in the tissue level and advance the understanding of mechanisms of baculovirus-induced host behavioral manipulation.

## Materials and Methods

### Insects, cultured cells, and viruses

*B. mori* larvae (Kinshu × Showa) were reared at 25°C with an artificial diet. BmN-4, Sf9, and Ld-652Y cells were maintained with TC100 medium supplemented with 10% fetal bovine serum. BmNPV T3 [42], AcMNPV C6 [43], and LdMNPV A21 [44] strains were used as wild-type viruses. Bm8D [29] was used as *Bm8* knockout BmNPV. BmNPV titers were determined using the plaque assay, and AcMNPV and LdMNPV titers were determined using the 50% tissue culture infectious dose (TCID_50_) method.

### Generation of recombinant viruses

Bm8OE, a *Bm8*-overexpressing virus, was generated as follows. The genomic region containing the *Bm8* coding sequence and its promoter was amplified from the genome of BmNPV T3 (GenBank accession no. L33180.1). The SV40 polyA sequence was amplified from the plasmid pIZ-GFP [45]. These two amplified fragments were inserted into the pBh-EPS vector [46] at the PstI site using the In-FusionHD Cloning Kit (Takara). The resultant plasmid and Bsu36I-digested BmNPV-abb genome DNA [47] were cotransfected into BmN-4 cells using FuGeneHD (Promega). The recombinant virus was purified twice using the plaque assay by identifying OB-producing plaques. *Bm8* overexpression was validated by quantitative polymerase chain reaction (qPCR).

Homologous proteins of Bm8 were searched by BLASTp against the nr database [48], and their coiled-coil domain was aligned. Point mutations of the conserved amino acid residues in the coiled-coil domain were introduced into a plasmid Bm8region-pcDNA [33] using a KOD-plus-Mutagenesis kit (Toyobo). The resultant plasmids were cotransfected with Bsu36I-digested Bm8D genome DNA, and the recombinant viruses were purified as described above by identifying white plaques.

All primers used are listed in Table S1.

### Locomotion assay

Locomotion assays were performed as reported previously [34]. Briefly, fourth-instar larvae were starved for several hours, injected intrahemocoelically with a viral suspension containing 10^5^ plaque-forming units (PFUs), and returned to an artificial diet at 25°C. At 48 hpi, each larva (n = 5) was placed in a 100 mm cell culture dish with a portion of an artificial diet. Locomotion was recorded at 3 s intervals using a time-lapse camera (TLC200-pro, Bruno) under continuous light conditions. The video was processed, and larval locomotory patterns and activities were analyzed using KaicoTracker, a recently developed software modified from the previous method [49].

### Viral virulence in infected larvae

Fifth-instar *B. mori* larvae were injected with a viral suspension as described above (n = 20). Dead and surviving larvae were counted from 96 hpi until all larvae died at 6 h intervals. The median lethal time (LT_50_) was determined using the Kaplan–Meier estimate.

### Western blotting

Fifth-instar *B. mori* larvae were injected with a viral suspension as described above. Hemolymph and fat bodies were collected at 90 hpi from three larvae and mixed into a single tube. Hemolymph was diluted with 2× sample buffer [100 mM Tris-HCl (pH6.8), 4% sodium deoxycholate, 12% 2-mercaptoethanol, 20% glycerol, and bromophenol blue]. Fat bodies were ground in a RIPA buffer [25 mM Tris-HCl (pH8.0), 150 mM NaCl, 1% NP-40, and 1% sodium deoxycholate] with 30 μg/ml E64 (Cayman Chemical), a cysteine protease inhibitor, followed by centrifugation to remove the insoluble fractions. The protein concentration of the supernatant was measured by Pierce BCA Protein Assay Kit (Thermo Fisher Scientific). The total protein amount was adjusted and diluted with a 2× sample buffer. A portion of the protein samples was subjected to Western blotting, as described previously [50].

### Cathepsin assay

The hemolymph of infected larvae was collected, and V-CATH activity was measured under the presence or absence of E64, as described previously (n =3) [50].

### Quantification of viral genomic DNA

BmN-4 cells were infected with BmNPV at a multiplicity of infection (MOI) of 5 and harvested at 0, 6, 12, 24, 48, and 72 hpi. The total DNA was extracted, and the amount of viral DNA was measured as described previously [51].

### Quantification of viral gene expression

The total RNA of BmNPV-infected BmN-4 cells was prepared with TRI reagent (Cosmo Bio Co., Ltd.) according to the manufacturer’s protocol. First-strand cDNA was synthesized from 1 μg total RNA using SuperScript® IV (Thermo Fisher Scientific), and qPCR was performed using the KAPA™ SYBR FAST qPCR kit (Kapa Biosystems) with the primers listed in Table S1. The expression level was calculated using the 2^-ΔΔCt^ method and represented as a relative expression when the expression in T3-infected cells at 72 hpi is 100.

### RNA sequencing

BmNPV-infected BmN-4 cells were harvested at 48 hpi, and total RNA was extracted as described above. Library preparation and sequencing were performed by Novagen Co., Ltd (China), using Novaseq6000 with 150 bp paired-end mode. The quality of sequence data was examined with FastQC. The reads were mapped to the BmNPV genome (GenBank accession no. L33180.1) using Hisat2 [52]. Resultant SAM files were converted to BAM files using SAMtools and subsequently converted to BED files using BEDTools [53, 54]. Viral gene expression was normalized to reads per kilobase of transcript per million, using mapped viral reads as the total number of reads. RNA sequencing data are available under accession number DRA013586.

Sequencing was performed using three biological replicates, but third datasets were clustered by the replicate rather than the treatments (Fig. S3A). Because the other two replicates were clustered by virus treatment (Fig. S3B), differential expression analysis was performed using the first and second replicates using DESeq2 [55], and the gene expression pattern was independently verified by the third replicate (Fig. S3C).

### Microscopic observation

Fifth-instar *B. mori* larvae were injected with a viral suspension as described above. MSGs were dissected from infected larvae at 90 hpi and washed in phosphate-buffered saline. The dissected tissues were observed under a stereomicroscope (Zeiss Axio Zoom.V16, Carl Zeiss).

### Transient expression of *bv/odv-e26*

*bv/odv-e26* homologus were amplified from the genomes of BmNPV T3, AcMNPV C6, and LdMNPV A21. The *gfp* coding region was amplified from the plasmid pIZ-GFP. These fragments were inserted into the pIZ-V5/His vector (Invitrogen) using the In-FusionHD Cloning Kit (Takara). Mutations in the coiled-coil domain were introduced by PCR with the primers listed in Table S1. The resultant plasmids were transfected into cultured cells susceptible to the original viruses using X-tremeGENE™ HP DNA transfection reagent (Roche). For BmNPV, cells were infected with the viruses at an MOI of 5 PFUs at 2 days post transfection (dpt). For AcMNPV and LdMNPV, cells were infected at an MOI of 5 TCID_50_ at 1 dpt. Expression levels of *ie1* and *polh* were measured by real-time qPCR as described above. Cells were observed at the designated time points (Figs. 3D and 4C and F) under a fluorescence microscope (FLoid Cell Imaging Station, Thermo Fischer Scientific).

### Homology searchs

ORFs located between *egt* and *Bm9* homologues were selected from the genomes of alphabaculoviruses registered in the NCBI RefSeq database. Their homologous genes were searched against alphabaculovirus genomes using the BLASTp [48]. Homologous relationships were defined by the reciprocal best hit. Network analysis was further performed, where genes and homologous relationships consist of nodes and edges, respectively (Fig. S6). Genes connected to *bv/odv-e26* in BmNPV were considered as *bv/odv-e26* homologues. All protein sequences of *bv/odv-e26* homologues were aligned with mafft version 7.453 [56]. Phylogenetic trees were constructed using IQtree [57] multicore version 1.6.12 under the LG model and the JTT+F+I+G4 model for concatenated core gene products and BV/ODV-E26, respectively, with 1000 ultrafast bootstraps. The constructed trees were visualized using iTOL [58]. The dataset of concatenated sequences of baculovirus core genes was retrieved from Harrison et al. [9].

### Statistical analysis

Locomotory distance, speed, and duration were compared using the log-rank test, followed by Bonferroni correction. The host survival time was compared using the pairwise log-rank test. V-CATH activity between treatments with DMSO and E64 was compared by the one-sided paired-samples *t*-test. Viral genome replication and temporal gene expression were compared using Tukey’s HSD test. In the transient expression experiment, viral gene expression was compared using the paired-samples *t*-test, and *p*-values were corrected to *q*-values using the Benjamini–Hochberg method.

## Acknowledgement

We thank Takashi Kiuchi, Munetaka Kawamoto, and Toru Shimada for clerical assistance. LdMNPV A21 and AcMNPV C6 strain were kindly provided from Dr. Motoko Ikeda at Nagoya University and Dr. Masashi Iwanaga at Utsunomiya University, respectively.

## Notes

### Competing Interest Statement

The authors have declared no competing interest.

