## Supplementally Figures for "Baculovirus *bv/odv-e26* is required for host behavioral manipulation by optimizing viral virulence to lepidopteran hosts"

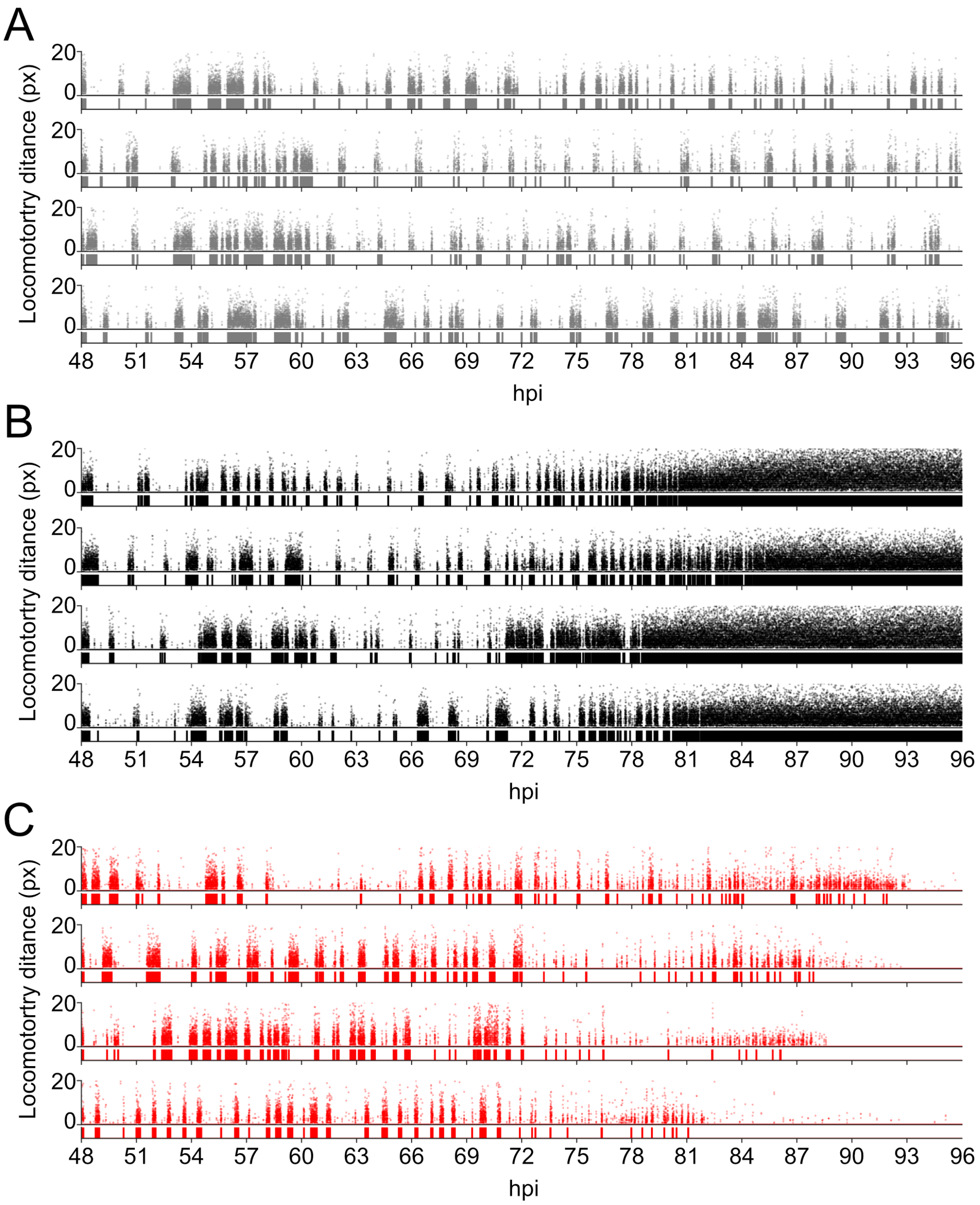


**Fig S1.** Behavioral patterns of the rest of the individuals that are not shown in Fig. 1. (A) Mock-infected, (B) T3-infected, and (C) Bm8D-infected larvae.

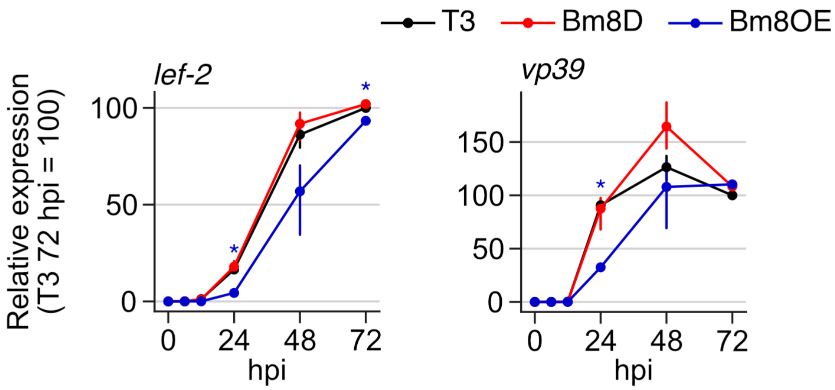


**Fig S2.** Temporal expression pattern of *lef-2* and *vp39*. Blue asterisks indicate a significant difference in Bm8OE from the other two viruses. $p < 0.05$, Tukey's HSD test. Error bars indicate 95% confidence intervals.

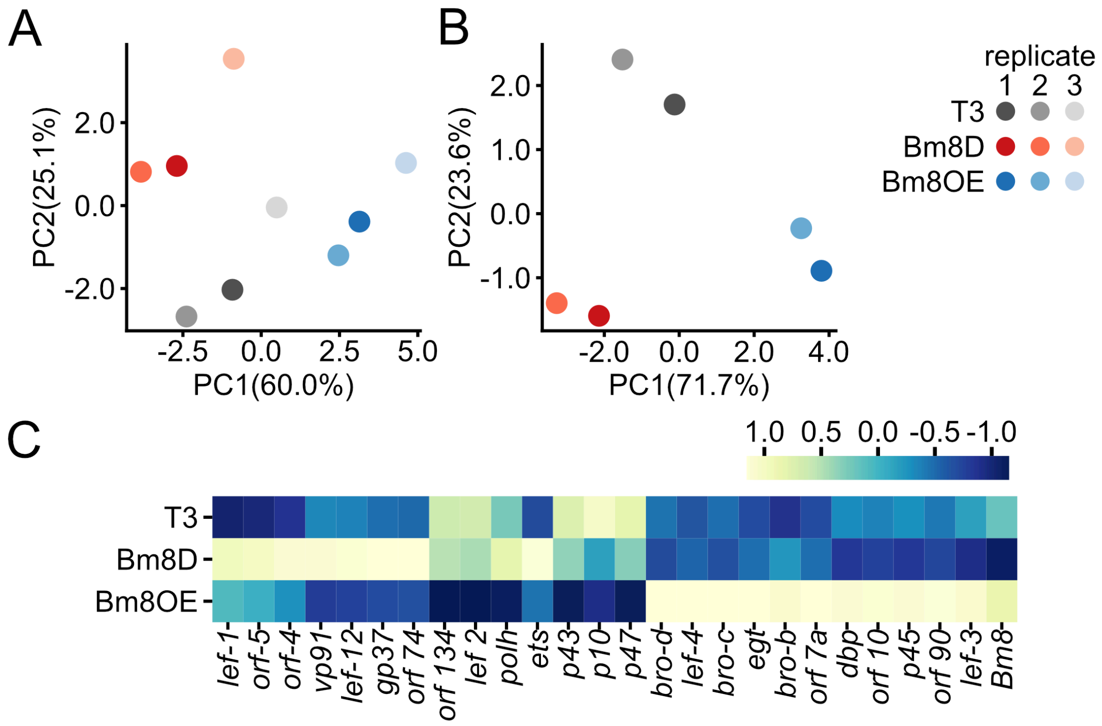


**Fig S3.** (A) Principal component analysis (PCA) results using three replicates. (B) PCA results using replicates 1 and 2. (A and B) Color circles indicate individual replicates. The percentage shown on each axis indicates the contribution of each principal component. (C) Heat map of viral gene expression in replicate 3. The order of genes corresponds to those in Fig. 2G.

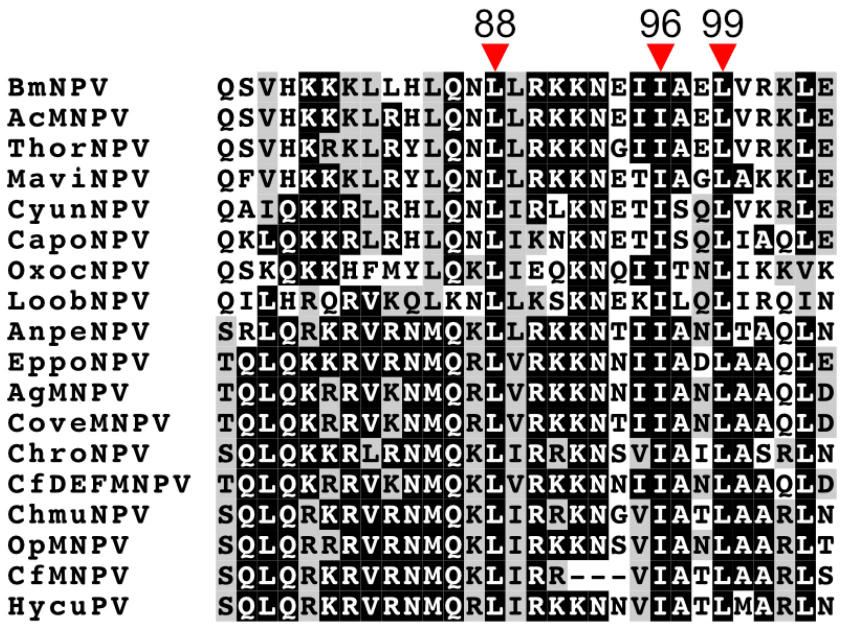


**Fig S4.** Amino acid sequence alignment of the coiled-coil domain of *bv/odv-e26* homologues. All viruses listed belong to group I alphabaculovirus. The abbreviations of virus names are listed in Table S2.

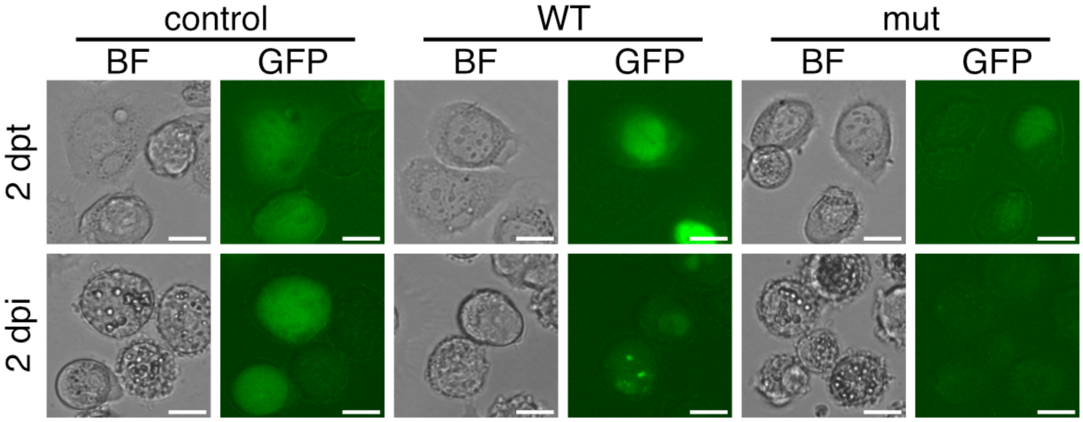


**Fig S5.** GFP fluorescence pattern in control-transfected, wild-type-transfected, and mutant-transfected BmN-4 cells at 2 dpt, followed by T3 infection at 2 dpi. Bars, 20 μm.

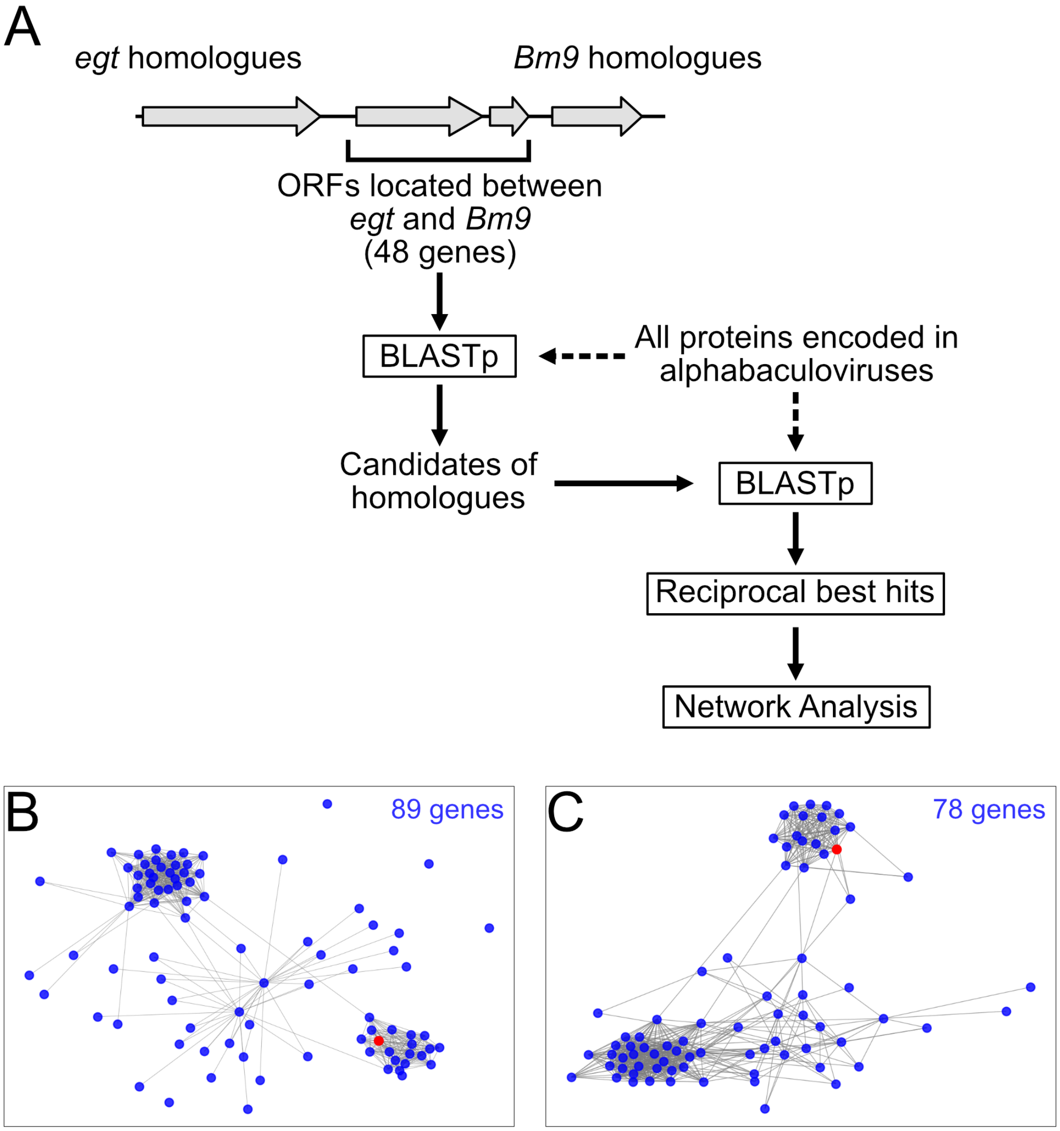


**Fig S6.** (A) Scheme of homology-based identification of *bv/odv-e26* homologues. (B) Network of homology relationship in 89 homologue candidates. (C) Network of the same relationship in 78 candidates showing the reciprocal best hits. (B and C) Each node and edge indicate gene and homology relationship, respectively. The red node represents *Bm8*.

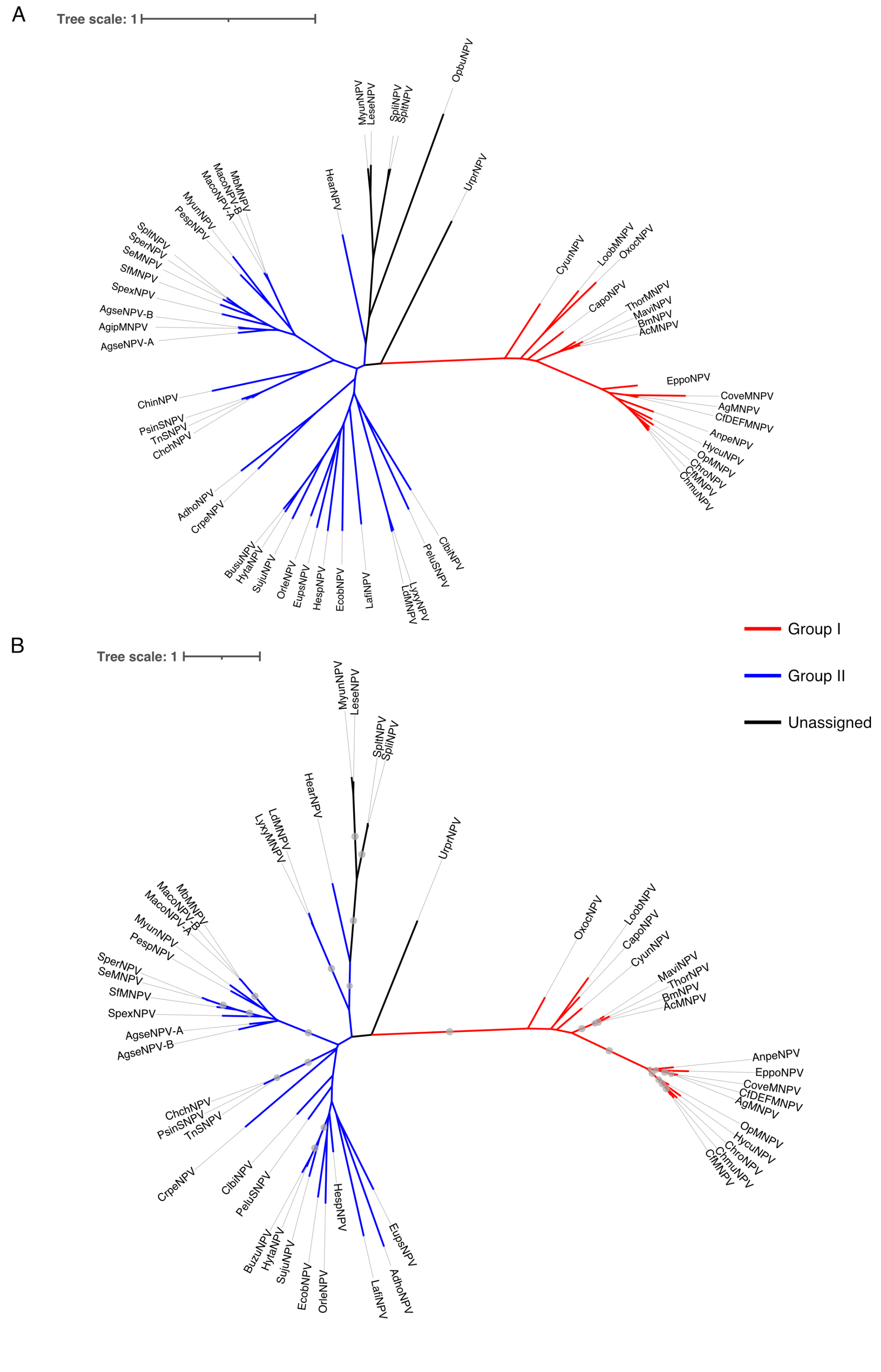


**Fig S7.** Unrooted phylogenic trees of (A) alphabaculoviruses, constructed based on 38 core genes with LG substitution model and (B) *bv/odv-e26* homologues, constructed under the JTT+F+I+G4 model with 1000 ultrafast bootstraps. Red, blue, and black indicate groups I and II and unassigned alphabaculoviruses, respectively. Gray circles indicate bootstrap values higher than 90. The abbreviations of virus names are listed in Table S2.

```
AdhoNPV      68  -NEKQTELIDKLKEEKERQENKIQLLRQSLV
AgipMNPV     79  YEQKYKIKVTELETALQRKCQKICQLQEDNP
AgseNPV-B    58  YEQKYKIKVSELETALQRKCQKIAQLQDDNT
AgseNPV-A    61  YEQKYKTKVVELEAALERKCRKIARLREESA
MbMNPV       54  YERKYKLKLVELEGALSHKKRQIEQLEEDIP
MacoNPV-B    54  YERKYKLKLVELEGALSHKKRQIEQLEEDIP
MacoNPV-A    54  YERKYKLKLVELEGALSHKKRQIEQLEEDIP
SperNPV      81  YERKYKSKVLELQTALQLKTQTIAQIEQDFV
SeMNPV       70  YERKYKSKVLELQTALQLKTQTIAQIEKDFV
SfMNPV        1  -------------------------------
SpexNPV      54  YEQKYKMKVFQLEVALDHKERKINKLHEDCR
MyunNPV      54  YDKKYKCKIDELQRAIDRKDEDIAELLEENS
PespNPV      54  YERKYKLKLANLQCSVETKRRRIAELQEKLS
AnpeNPV      60  -SRLQRKRVRNMQKLLRKKNTIIANLTAQLN
AgMNPV       73  -TQLQKRRVKNMQRLVRKKNNIIANLAAQLD
CfDEFMNPV    73  -TQLQKRRVKNMQKLVRKKNNIIANLAAQLD
CoveMNPV     73  -TQLQKRRVKNMQRLVRKKNTIIANLAAQLD
EppoNPV      63  -TQLQKKRVRNMQRLVRKKNNIIADLAAQLE
CfMNPV       61  -SQLQRKRVRNMQKLIRR---VIATLAARLS
ChmuNPV      57  -SQLQRKRVRNMQKLIRRKNGVIATLAARLN
ChroNPV      60  -SQLQKKRLRNMQKLIRRKNSVIAILASRLN
HycuNPV      60  -SQLQRKRVRNMQRLIRKKNNVIATLMARLN
OpMNPV       60  -SQLQRRRVRNMQKLIRKKNSVIANLAARLT
HycuNPV       2  -------------------------------
AcMNPV       75  -QSVHKKKLRHLQNLLRKKNEIIAELVRKLE
BmNPV        75  -QSVHKKKLLHLQNLLRKKNEIIAELVRKLE
ThorNPV      73  -QSVHKRKLRYLQNLLRKKNGIIAELVRKLE
MaviNPV      63  -QFVHKKKLRYLQNLLRKKNETIAGLAKKLE
CyunNPV      84  -QAIQKKRLRHLQNLIRLKNETISQLVKRLE
CapoNPV     112  -QKLQKKRLRHLQNLIKNKNETISQLIAQLE
LoobNPV      86  -QILHRQRVKQLKNLLKSKNEKILQLIRQIN
OxocNPV      46  -QSKQKKHFMYLQKLIEQKNQIITNLIKKVK
UrprNPV      59  CEEIYKNKICKLELQIEQKDCIIENLRHEIN
HearNPV      59  ----YRAQINILKKSLRHKQQIIDELKDKLS
LeseNPV     101  --------INMLKLALVNERKKYQTLLSEEG
MyunNPV     144  --------INMLKLALVNERKKYQSLLTDRK
SpliNPV      46  --QQQRQTIRLLRNTLKQK------FKNEIA
SpltNPV      46  --QQQRQTIRSLRNTLKQK------IKNEIT
LdMNPV       64  CNLQYKNRVRFLRRVLAKRTARVDRLRREHG
LyxyMNPV     63  CNLQYKNKVRFLRRVLAKRNARIDRLRREHG
LeseNPV      47  -------------GFIGEHVSRLSQLARNLL
ChchNPV      56  YAERYKNKIQNLQNTIDKKDRHIKDLYKQID
PsinSNPV     57  YAERYKNKIENLQNSIDKKDRHIKDLYKQID
TnSNPV       56  YAERYKNKIQNLQNSIDKKDRHIKDLYKQID
BuzuNPV      62  YEQKYEMRLERLRKLLCKKTRKLKILQKKFN
HytaNPV      59  YEQKYEMRLQRLRKVLSKRTRKLNTLQKRIN
SujuNPV      91  YKQRYEMQLQRLQTLLAKRTRKLQNLRQKLS
EcobNPV      70  YRQEYDARLQKLYVSLKRKTMKLEKLRRRKE
OrleNPV      76  YEHKYRSRFIKLRSLLSKKQRKIEHLKKINK
HespNPV      72  YERQYEVRLQRLRLLLGKKTRKLQRLKTKFN
EupsNPV      90  YKKRYEIRVKKLRLSLFKKLKKIQNFDKIAQ
LafiNPV     141  YRYECTARIKKMINLLEKKDRRIEKLKKRYK
PeluSNPV     58  YDDKYKIKIQRLRAIIAKKNRKIRSLRNECF
ClbiNPV      91  YEEKYKAKIQRLRADIAKKNRKITRLRNKIQ
CrpeNPV     104  DKHEFEISTEKLKNTIRRKDTKIKQLQSKVK
```

**Fig S8.** Alignment of the coiled-coil domain region of all *bv/odv-e26* homologues. HycuNPV has two separated proteins that correspond to the N- and C-terminal regions of the protein, respectively. LeseNPV have two homologues. The SfMNPV homologue lacks the corresponding region. The abbreviations of virus names are listed in Table S2.

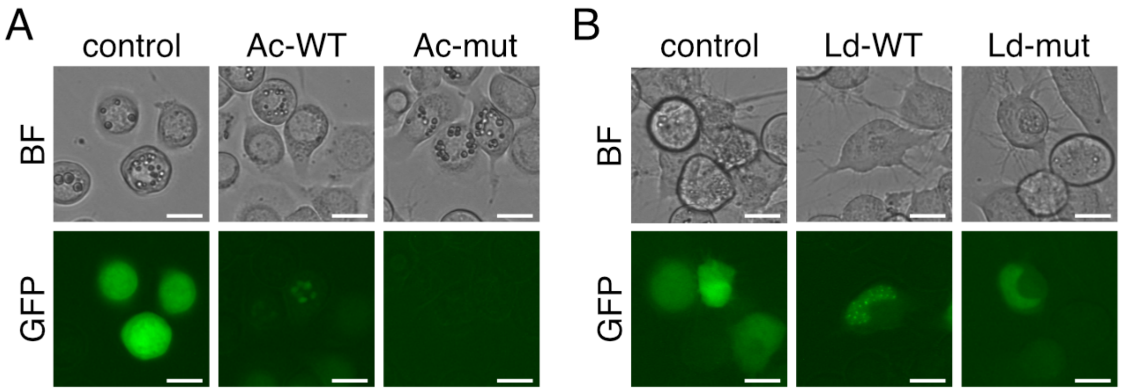


**Fig S9.** Localization of the *bv/odv-e26* homologues of AcMNPV and LdMNPV. (A) AcMNPV-infected Sf-9 and (B) LdMNPV-infected Ld652Y cells at 72 hpi. Bars, 20 µm.

**Table S1.** Primer list.

| Name | Use | Nucleotide Sequene (5′→3′) |
|---|---|---|
| Bm8-FLAG-F-iP | Generation of Bm8OE | GACGATGACAAGTAATTTGAAGGGTGAGGAAGAGCCCAATTG |
| Bm8-FLAG-R-iP | Generation of Bm8OE | ATCCTTGTAGTCATAGGCGTTAATATCACTTTGAGATTCATCTTGC |
| Bm8FLAG-F-IF | Generation of Bm8OE | ATTTGTATCGGAGCTCGATTTACGTAAGTTTTGGG |
| Bm8FLAG-R-IF | Generation of Bm8OE | CTGCAATAAACAAGTTTACTTGTCATCGTCATCC |
| sv40pA-F-IF | Generation of Bm8OE | ACTTGTTTATTGCAGCTTATAATG |
| sv40pA-R-IF | Generation of Bm8OE | ATAAATGTCAGAATTACAGACATGATAAGATACATTGATGAG |
| Bm8regionF | Sequence confrimation | CTCGAGGAACCTACATCCCATATTTGACAAC |
| Bm8regionR | Sequence confrimation | AAGCTTGTACAATTCGTGTGTCAATGATCTC |
| Bm8L88A-F | Generation of Bm8-L88A | CTAAGGAAAAAGAACGAAATTATTGC |
| Bm8L88A-R | Generation of Bm8-L88A | CGCATTTTGCAAATGCAGCAGTTTC |
| Bm8I96A/L99A-F | Generation of Bm8-I96A and Bm8-L99A | GTTAGAAAACTCGAAAGTGCAC |
| Bm8I96A-R | Generation of Bm8-I96A | CAACTCGGCAGCAATTTCGTTCTTTTTCC |
| Bm8L99A-R | Generation of Bm8-L99A | CGCCTCGGCAATAATTTCGTTCTTTTTCC |
| GFPgene-EcoRI-Fn | generaion of GFP-Bm8 | ATCGAATTCTAAGGTGAGCAAGGGCGAGGAGCTG |
| GFPgene-EcoRI-Rnn | generaion of GFP-Bm8 | AGTGAATTCGCACACTTGTACAGCTCGTCCATGCC |
| GFP-L-Bm8-iF | Inserting linker sequence between *gfp* and *Bm8* | CACCGCCACTTCCTCCACCGCCCGCACACTTGTACAGCTCGTCCATG |
| GFP-L-Bm8-iR | Inserting linker sequence between *gfp* and *Bm8* | GAGGTTCAGGTGGTGGCGGATCGAATTCTGTTCACACGCGCTTATGTG |
| Bm8F | Construction of pIZ-Bm8LG | GTACCGAGCTCGGATCCATGAATTCTGTTCACACGCG |
| Bm8-gfpR | Construction of pIZ-Bm8LG | ACTGCCTCCACCGCCAGAGCCACCTCCACCATAGGCGTTAATATCACTTTG |
| gfpF | Construction of pIZ-Bm8LG | GGCGGTGGAGGCAGTGGAGGTGGCGGATCGTTGGTGAGCAAGGGCGAGGAG |
| gfpR | Construction of pIZ-Bm8LG | CTGGACTAGTGGATCCTCACTTGTACAGCTCGTCCATGCC |
| Bm8/Ac16-gfpR | Construction of pIZ-Bm8LG, pIZ-Ac16LG | ACTGCCTCCACCGCCAGAGCCACCTCCACCATAGGCGTTAATATCACTTTG |
| Ac16F | Construction of pIZ-Ac16LG | GTACCGAGCTCGGATCCATGGAGTCTGTTCAAACGCG |
| Ld127F | Construction of pIZ-Ld127 | GTACCGAGCTCGGATCCATGTCGACTTGGAGGAACAAATTGC |
| Ld127-gfpR | Construction of pIZ-Ld127 | ACTGCCTCCACCGCCAGAGCCACCTCCACCGTGCCGCCGGCGGCG |
| Bm8CCmutF | Introducing mutaitons in coiled-coil domain | GAACGAAATTGCTGCCGAGGCGGTTAGAAAACTC |
| Bm8CCmutR | Introducing mutaitons in coiled-coil domain | GAGTTTTCTAACCGCCTCGGCAGCAATTTCGTTC |
| Ac16CCmutF | Introducing mutaitons in coiled-coil domain | GAAAAAAGAACGAAATTGCTGCCGAGGCGGTTAGAAAACTTG |
| Ac16CCmutR | Introducing mutaitons in coiled-coil domain | CAAGTTTTCTAACCGCCTCGGCAGCAATTTCGTTCTTTTTTC |
| Ld127CCmutF | Introducing mutaitons in coiled-coil domain | GGACCGCTCGCGCAGACCGAGCGCGCCGCGAGCAC |
| Ld127CCmutR | Introducing mutaitons in coiled-coil domain | GTGCTCGCGGCGCGCTCGGTCTGCGCGAGCGGTCC |
| ago3_gF | Quantification of host genome | TTTCTTAGTACACTCAAACG |
| ago3_gR | Quantification of host genome | CTCTCTTCGTAGAACATATC |
| polhqF | Quantification of *polh* and viral genome | GAACAAGAGGAGAAGCAATG |
| polhqR | Quantification of *polh* and viral genome | TCCAGTTGGCGATTAACTTC |
| ie1qF | Quantification of *ie1* | TACTTGGACGATTCACAAAG |
| ie1qR | Quantification of *ie1* | GTGCAAATGTTCGTGTTGTG |
| lef2qF | Quantification of *lef2* | ACATGCTGAACAGCAAGATC |
| lef2qR | Quantification of *lef2* | ACATCGGTTTTCACATTTGG |
| vp39qF | Quantification of *vp39* | ACTTTTCATGATGTCACTGC |
| vp39qR | Quantification of *vp39* | AGTACTTGCAAATCGACACG |
| Bm8qF | Quantification of *Bm8* | AACTCGAAAGTGCACAGAAG |
| Bm8qR | Quantification of *Bm8* | CAATAATTGTGCGAATTGTG |
| Ac_ie1qF | Quantification of AcMNPV *ie1* | TCACGTACAAATACAGCAGCGTCG |
| Ac_ie1qR | Quantification of AcMNPV *ie1* | CATGCTGCCGTCCTCCTTCTTAAC |
| Ac_polhqF | Quantification of AcMNPV *polh* | GTTACAAATTCCTGGCCCAACAC |
| Ac_polhqR | Quantification of AcMNPV *polh* | ATGCGGTACTCGTTGTTGCTG |
| Ac16qF | Quantification of *Ac16* | TGCACAGAAGAAGACAACGCAC |
| Ac16qR | Quantification of *Ac16* | CGGCCAAACGTCTCCTTACAAAC |
| Ld_ie1qF | Quantification of LdMNPV *ie1* | CCCGTCGAACTTGTTGATGATGTC |
| Ld_ie1qR | Quantification of LdMNPV *ie1* | AAGAACGAGGAGCGCCTGAC |
| Ld_polhqF | Quantification of LdMNPV *polh* | AAAGCACTTGGAACAGCACGAG |
| Ld_polhqR | Quantification of LdMNPV *polh* | GGCTTGACATTGCGGATCTCTTTG |
| Ld127qF | Quantification of *Ld127* | ATTGGCGCGTGATTTCGGTG |
| Ld127qR | Quantification of *Ld127* | ATATTGTAGGTTGCACCGCTCC |

**Table S2.** List of alphabaculovirus genomes used in this study.

| Species | Isolate | Accession number | Virus Abbrev. |
|---|---|---|---|
| *Adoxophyes honmai nucleopolyhedrovirus* | ADN001 | AP006270 | AdhoNPV |
| *Agrotis ipsilon multiple nucleopolyhedrovirus* | Illinois | EU839994 | AgipMNPV |
| *Agrotis segetum nucleopolyhedrovirus A* | Polish | DQ123841 | AgseNPV-A |
| *Agrotis segetum nucleopolyhedrovirus B* | English | KM102981 | AgseNPV-B |
| *Antheraea pernyi nucleopolyhedrovirus* | Liaoning | DQ486030 | AnpeNPV |
| *Anticarsia gemmatalis multiple nucleopolyhedrovirus* | 2D | DQ813662 | AgMNPV |
| *Autographa californica multiple nucleopolyhedrovirus* | C6 | L22858 | AcMNPV |
| *Bombyx mori nucleopolyhedrovirus* | T3 | L33180 | BmNPV |
| *Buzura suppressaria nucleopolyhedrovirus* | Hubei | KF611977 | BuzuNPV |
| *Catopsilia pomona nucleopolyhedrovirus* | 416 | KU565883 | CapoNPV |
| *Choristoneura fumiferana DEF multiple nucleopolyhedrovirus* | | AY327402 | CfDEFMNPV |
| *Choristoneura fumiferana multiple nucleopolyhedrovirus* | Ireland | AF512031 | CfMNPV |
| *Choristoneura murinana nucleopolyhedrovirus* | Darmstadt | KF894742 | ChmuNPV |
| *Choristoneura rosaceana nucleopolyhedrovirus* | NB_1 | KC961304 | ChroNPV |
| *Chrysodeixis chalcites nucleopolyhedrovirus* | | AY864330 | ChchNPV |
| *Chrysodeixis includens nucleopolyhedrovirus* | IE | KJ631622 | PsinSNPV |
| *Clanis bilineata nucleopolyhedrovirus* | DZ1 | DQ504428 | ClbiNPV |
| *Condylorrhiza vestigialis nucleopolyhedrovirus* | PR.2002 | KJ631623 | CoveMNPV |
| *Cryptophlebia peltastica nucleopolyhedrovirus* | SA | MH394321 | CrpeNPV |
| *Cyclophragma undans nucleopolyhedrovirus* | Whiov | KT957089 | CyunNPV |
| *Ectropis obliqua nucleopolyhedrovirus* | A1 | DQ837165 | EcobNPV |
| *Epiphyas postvittana nucleopolyhedrovirus* | | AY043265 | EppoNPV |
| *Euproctis pseudoconspersa nucleopolyhedrovirus* | Hangzhou | FJ227128 | EupsNPV |
| *Helicoverpa armigera nucleopolyhedrovirus* | G4 | AF271059 | HearNPV |
| *Hemileuca species nucleopolyhedrovirus* | MEM | KF158713 | HespNPV |
| *Hyphantria cunea nucleopolyhedrovirus* | N9 | AP009046 | HycuNPV |
| *Hyposidra talaca nucleopolyhedrovirus* | India.001 | MH261376 | HytaNPV |
| *Lambdina fiscellaria nucleopolyhedrovirus* | GR15 | KP752043 | LafiNPV |
| *Leucania separata nucleopolyhedrovirus* | AH1 | AY394490 | LeseNPV |
| *Lonomia obliqua nucleopolyhedrovirus* | SP/2000 | KP763670 | LoobNPV |
| *Lymantria dispar multiple nucleopolyhedrovirus* | 5-6 | AF081810 | LdMNPV |
| *Lymantria xylina nucleopolyhedrovirus* | 5 | GQ202541 | LyxyMNPV |
| *Mamestra brassicae multiple nucleopolyhedrovirus* | K1 | JQ798165 | MbMNPV |
| *Mamestra configurata nucleopolyhedrovirus A* | 90/2 | U59461 | MacoNPV-A |
| *Mamestra configurata nucleopolyhedrovirus B* | 96B | AY126275 | MacoNPV-B |
| *Maruca vitrata nucleopolyhedrovirus* | MV-8 | EF125867 | MaviNPV |
| *Mythimna unipuncta nucleopolyhedrovirus A* | #7 | MF375894 | MyunNPV |
| *Mythimna unipuncta nucleopolyhedrovirus B* | KY310 | MH124167 | MyunNPV |
| *Operophtera brumata nucleopolyhedrovirus* | MA | MF614691 | OpbuNPV |
| *Orgyia leucostigma nucleopolyhedrovirus* | CFS-77 | EU309041 | OrleNPV |
| *Orgyia pseudotsugata multiple nucleopolyhedrovirus* | | U75930 | OpMNPV |
| *Oxyplax ochracea nucleopolyhedrovirus* | 435 | MF143631 | OxocNPV |
| *Peridroma saucia nucleopolyhedrovirus* | GR167 | KM009991 | PespNPV |
| *Perigonia lusca nucleopolyhedrovirus* | | KM596836 | PeluSNPV |
| *Spodoptera eridania nucleopolyhedrovirus* | 251 | MH320559 | SperNPV |
| *Spodoptera exempta nucleopolyhedrovirus* | 244.1 | MH717816 | SpexNPV |
| *Spodoptera exigua multiple nucleopolyhedrovirus* | US1 | AF169823 | SeMNPV |
| *Spodoptera frugiperda multiple nucleopolyhedrovirus* | 3AP2 | EF035042 | SfMNPV |
| *Spodoptera littoralis nucleopolyhedrovirus* | AN1956 | JX454574 | SpliNPV |
| *Spodoptera litura nucleopolyhedrovirus* | G2 | AF325155 | SpltNPV |
| *Sucra jujuba nucleopolyhedrovirus* | 473 | KJ676450 | SujuNPV |
| *Thysanoplusia orichalcea nucleopolyhedrovirus* | p2 | JX467702 | ThorNPV |
| *Trichoplusia ni single nucleopolyhedrovirus* | | DQ017380 | TnSNPV |
| *Urbanus proteus nucleopolyhedrovirus* | Southern Brazil | KR011717 | UrprNPV |
